# Alpha blocking and 1/*f^β^* spectral scaling in resting EEG can be accounted for by a sum of damped alpha band oscillatory processes

**DOI:** 10.1101/2021.08.20.457060

**Authors:** Rick Evertz, Damien G. Hicks, David T. J. Liley

## Abstract

The dynamical and physiological basis of alpha band activity and 1/*f^β^* noise in the EEG are the subject of continued speculation. Here we conjecture, on the basis of empirical data analysis, that both of these features may be economically accounted for through a single process if the resting EEG is conceived of being the sum of multiple stochastically perturbed alpha band damped linear oscillators with a distribution of dampings (relaxation rates). The modulation of alpha-band and 1/*f^β^* noise activity by changes in damping is explored in eyes closed (EC) and eyes open (EO) resting state EEG. We aim to estimate the distribution of dampings by solving an inverse problem applied to EEG power spectra. The characteristics of the damping distribution are examined across subjects, sensors and recording condition (EC/EO). We find that there are robust changes in the damping distribution between EC and EO recording conditions across participants. The estimated damping distributions are found to be predominantly bimodal, with the number and position of the modes related to the sharpness of the alpha resonance and the scaling (*β*) of the power spectrum (1/*f^β^*). The results suggest that there exists an intimate relationship between resting state alpha activity and 1/*f^β^* noise with changes in both governed by changes to the damping of the underlying alpha oscillatory processes. In particular, alpha-blocking is observed to be the result of the most weakly damped distribution mode becoming more heavily damped. The results suggest a novel way of characterizing resting EEG power spectra and provides new insight into the central role that damped alpha-band activity may play in characterising the spatio-temporal features of resting state EEG.

**Author summary:** The resting human electroencephalogram (EEG) exhibits two dominant spectral features: the alpha rhythm (8-13 Hz) and its associated attenuation between eyes-closed and eyes-open resting state (alpha blocking), and the 1/*f^β^* scaling of the power spectrum. While these phenomena are well studied a thorough understanding of their respective generative processes remains elusive. By employing a theoretical approach that follows from neural population models of EEG we demonstrate that it is possible to economically account for both of these phenomena using a singular mechanistic framework: resting EEG is assumed to arise from the summed activity of multiple uncorrelated, stochastically driven, damped alpha band linear oscillatory processes having a distribution of relaxation rates or dampings. By numerically estimating these damping distributions from eyes-closed and eyes-open EEG data, in a total of 136 participants, it is found that such damping distributions are predominantly bimodal in shape. The most weakly damped mode is found to account for alpha band power, with alpha blocking being driven by an increase in the damping of this weakly damped mode, whereas the second, and more heavily damped mode, is able to explain 1/*f^β^* scaling present in the resting state EEG spectra.

## Introduction

Electroencephalography (EEG) is a non-invasive method used to measure the electrical activity of the brain at the surface of the scalp, with the recorded voltage fluctuations being generated by ionic current flows resulting from neural synaptic activity across the cortex [1]. EEG time series recordings display a rich array of dynamic activity that is distributed spatio-temporally across the head, providing a unique window into the inner workings of the human brain. Since its discovery the EEG has been widely used as a sensitive measure of brain state in disease and health [2,3]. We aim to explore two prominent features of the EEG that are well observed but of which the mechanistic origins are the subject of continued speculation; the alpha rhythm and its changes between eyes-closed (EC) and eyes-open (EO) resting states, and the 1/*f^β^* frequency scaling of the power spectrum.

First discovered and recorded in the early 20th century by Hans Berger [4], the alpha rhythm (8-13 Hz), also commonly known as alpha band activity, is arguably the most dominant feature of the resting EEG. There exists significant heterogeneity of alpha band activity across the population in the terms of its the magnitude of its activity, its peak/central frequency and its reactivity to changes in behavioural and physiological state [5]. Topographically alpha band activity can be recorded across the scalp with it being particularly prominent occipitally [6]. Despite nearly a century of detailed empirical investigation, the physiological mechanisms responsible for the genesis of alpha rhythm remain essentially unresolved. The prevailing view is that the thalamus is central to the generation of the alpha rhythm through reverberant feed forward and feedback interactions between thalamic and cortical neuron populations [7,8]. Such a view was born out of earlier conceptions where alpha oscillations, intrinsically generated in the thalamus were thought to directly drive activity in overlying cortex [9]. However, other attempts to explain the genesis of the alpha rhythm also exist that depend on a mean field or neuronal population framework to model reverberant activity solely between excitatory and inhibitory cortical neuronal populations [10]. Complicating efforts to develop mechanistically and physiologically coherent accounts of the genesis of alpha band activity is the well described reduction in alpha band power between EC and EO conditions and in response to the exertion of mental effort, known commonly as alpha blocking [8, 11].

Alpha blocking is most obvious in the power spectral density of the recorded resting EEG, where a significant attenuation of peak alpha power is observed in the transition from EC to EO states. Such changes in power are also observed across a range of cognitive tasks [12,13]. The current view is that alpha blocking is thought to be caused by changes to the synchronous activity of neural populations across the cortex, where reductions in phase synchrony of neural activity are directly responsible for reductions in alpha peak power [14]. For this reason event related inreases/decreases in alpha band activity are often mechanistically designated as Event Related Synchronization/Event Related Desynchronization (ERD/ERS).

Another prominent feature of the resting EEG is the 1/*f^β^* (*β* ≈ 1 – 2) scaling of the power spectral density [28]. Such scaling, often referred to a ‘1/*f*’ noise, is also apparent in the power spectral densities of a range of time varying systems [15,16]. Because such power law scaling is independent of frequency, and thus the temporal scale, the corresponding dynamical activity is often referred to as ‘scale free’ [17]. ‘1/*f*’ noise has been observed in most forms of recorded brain activity, including electrocortigraphy, blood oxygen level dependent functional magnetic resonance and the magnetoencephalogram (MEG) [18–20]. The received view is that rhythmic activity such as the alpha rhythm occurs on a background of arrhythmic ‘1/*f*’ noise. In general the quantitative analysis of the EEG involves the decomposition into the dominant frequency bands, in attempt to find band power changes that correlate with cognition or behavioral states [21], and disregards the arrhythmic ‘1/*f*’ component as not being physiologically or behaviourally important. However, in recent years the functional relevance of ‘1/*f*’ noise has been reevaluated with it been suggested that such activity may play a role not only in healthy brains, but also in disease and psychological disorder [21–23]. For example age related changes in ‘1/*f*’ noise have been documented where the power spectra becomes more ‘white’ (reduced power exponent) with increasing age [24], suggesting that a decline in mental faculties may be due to more noisy neural activity.

Despite significant empirical investigations into the characterisation of such ‘1/*f*’ noise in brain activity, the dynamical and physiological mechanisms responsible for its generation in the brain remain unclear. The prevailing view hypothesizes that such scale free activity is the consequence of critical dynamics, where the brain is self organizing to a region of near-criticality thought to be the optimal for information transmission [25]. However, whether the brain exhibits self organized criticality remains controversial [26]. For this reason alternative explanations have been sought, that include the speculation that ‘1/*f*’ noise arises a consequence of the low-pass filtering of the local field potential by the neuronal dendrites of cortical excitatory neurons [27].

Nevertheless, despite such neurophysiological mechanistic uncertainty general models for the generation of ‘1/*f*’ noise do exist. For example it is well known that 1/*f* spectral profiles can be modelled as the the sum of a population of damped linear oscillators having a uniform distribution of dampings (relaxation rates) [16]. Such an approach has been used to explain the relation between alpha band activity and ‘1/*f*’ noise that is measured when investigating the pharmacologically induced alterations to MEG spectral scaling [28], and thus suggests alternative mechanisms to criticality for the genesis of scale free activity: ‘1/*f*’ noise could arise simply from a mixture of alpha band relaxation processes that having a distribution of dampings.

To date efforts to model the dynamic activity of the EEG fall into two broad categories. The first and most intuitive is the network modeling approach, which attempts to construct the EEG dynamics from the ground up by focusing on modelling individual cortical neurons and their vast networks of interactions. Such methods are limited by the computational uncertainty of simulating the interaction of hundreds of millions of neurons that lie under even a single EEG sensor. The ambiguity of how much physiological detail to include in these network approaches and how to meaningfully parameterize the model is a critical challenge [29]. An alternative approach are neural population models or mean-field theories. Motivated by the fact that a single EEG electrode records the aggregate activity of millions of cortical neurons, mean-field models account for EEG activity by modelling the average activity of interconnected populations of excitatory and inhibitory neurons in cortex and thalamus [30]. Mean-field models treat cortical and thalamic tissue as spatially continuous and model mesoscopic-scale cortical dynamics, and thus provide a major advantage in that they are constructed at spatial scales that are more commensurate with the physical coverage of an individual EEG channel (mm^2^ - cm^2^) [29]. Given that most mean field approaches generally divide the surface of the brain into populations of interconnected cortical neuron populations and investigates bulk dynamic properties, the number of modelled elements is drastically fewer than a neural network approach.

Mean field models can be mathematically quite compact and are typically formulated as multiple coupled nonlinear ordinary or partial differential equations [29]. Under suitable assumptions the defining non-linear equations can be linearised and in doing so reveal a rich range of electroencepahlographically plausible features including alpha band activity [10,31,32]. In such linearisations alpha band oscillatory activity is generally accounted through a linear time invariant transfer function driven by broad-band white noise, which pragmatically accounts for the spatio-temporal complexity of extra-cortical neuronal input. This approach is generally agnostic to the specific mean-field model used and enables the derivation of an electrocortical transfer function that describes the spectral response of the system to white noise driving. Given that most mean-field models of the EEG attempt to produce the alpha rhythm while simultaneously requiring a stable linearisation, the general form of the electrocortical impulse response embodies two essential features: i) it will produce oscillations that fall within the alpha band, and ii) will decay away to zero after perturbation, ensuring its stability. Regardless of the particular mean field theory chosen, qualitatively the modelled spectral behaviour of the resting EEG will be similar in form; a sharp alpha resonance followed by decline in spectral power with frequency.

The current uncertainty associated with the mechanisms responsible for alpha blocking and ‘1/*f*’ noise in resting state EEG leaves many open questions. For this reason we explore whether an alternative mechanism, that is theoretically motivated by mean-field models of the EEG and a general model for 1/*f* noise, can account for both of these phenomena within a single mechanistic framework. To achieve this we construct a simple electrocortical impulse response that approximates the main spectral features of the recorded EEG signal. On this basis the simplest electrocortical impulse response for the system under investigation is a damped linear oscillation (specifically a decaying cosinusoid) that is parameterised by its oscillation frequency (*f_α_* = 8 – 13 Hz) and damping (*γ*). In the frequency domain the power spectral density of such an impulse response will appear approximately Lorentzian in shape, in that it will have a sharp alpha peak and scale inversely with the square of the frequency (see Fig 1a). By extending this description to a more physiologically plausible case where the EEG signal recorded at a single location is composed of a sum of damped linear oscillations arising from multiple neuronal populations, we can take advantage of a general approach in which 1/*f^β^* power spectral profiles can arise as the result of a sum of exponentially decaying processes having a uniform distribution of decay rates [16] (Fig 1b). Therefore, we hypothesise that the resting EEG power spectrum can be well described as a superposition of uncorrelated damped alpha band linear oscillations that have a distribution of dampings. On this basis we seek to determine whether changes in the underlying distribution of dampings of alpha band relaxation processes are able to sufficiently account for ‘1/*f*’ noise and the spectral changes associated with alpha blocking, using two freely available EEG data sets (combined total of 136 participants) containing EC and EO resting state time series data. The empirical estimation of the distribution of dampings are found by solving an inverse problem by applying regularization methods to a defining Fredholm integral equation of the first kind.

**Fig 1.**
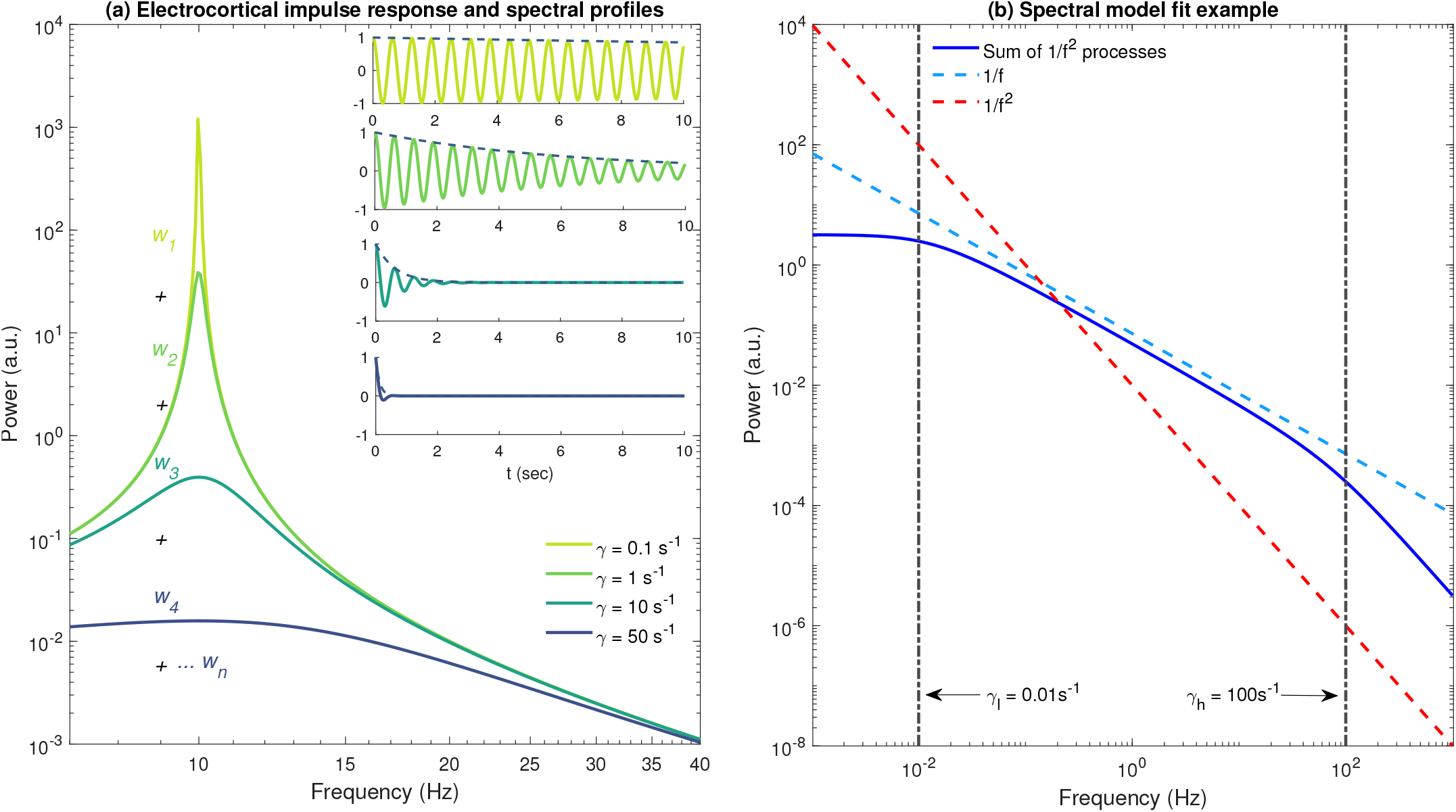
Resting EEG as a sum of damped independent alpha oscillatory processes. **(a)** Power spectral density of the model electrocortical impulse response cos(2*πf_α_t*) exp(−*γt*)Θ(*t*) for *γ* = {0.1,1,10, 50} *s*^−1^, where Θ(*t*) is the Heaviside step function. Resting EEG is modeled as a continuous sum of such processes over some suitable interval of dampings [*γ_l_, γ_h_*]. In general we aim to find the appropriate weights {*w_i_*} that account for the shape of the resting EEG power spectrum. **(b)** A 1/*f* spectrum (dashed blue) can be approximated as a sum of 1/*f*^2^ relaxation processes (solid blue) that are uniformly distributed in damping over some interval which for the displayed case is [*γ_l_* = 10^−2^, *γ_h_* = 10^2^] s^−1^. For clarity we have assumed *f_α_* =0 Hz.

The paper is organized as follows. The method section begins with a detailed mathematical specification of the inverse problem to be solved in order to estimate the distribution of alpha relaxation oscillation dampings from recorded EEG power spectral densities. This is then followed by a description of the Tikhonov regularisation method used to numerically solve the the corresponding Fredholm integral equation of the first kind. Finally in the methods, we detail the empirical EEG data sets used and the procedures used to ensure the numerical fidelity, and the properties, of our inverse solutions. Detailed results are then presented, followed by a discussion regarding the essential results.

## Results

### Numerical validation of Tikhonov regularised inverse method

To ensure that our inverse method was able to meaningfully recover damping probability distributions we conducted a simulation study. Six distinct prior damping distributions (Fig 2a-f) together with 100 regularised parameters spanning the interval [10^−6^ ≤ λ ≤ 1] were generated. The damping distributions for a given regularisation parameter that resulted in the minimum RSS error was deemed the best solutions. The regularisation scheme performed well for a unimodal distributions (Fig 2a). For bimodal distributions, the regularisation method correctly recovered the number and position of modes for both cases (equal and different peak magnitudes) with small discrepancies evident in the second mode where the peak amplitude was less in the regularised damping distribution than in the test distribution (Fig 2b-c). The trimodal distribution with constant peak amplitude (Fig 2d) was recovered well with only minor variations recorded in the width of the second and third mode. The recovery for the trimodal distributions with variable mode amplitudes (Fig 2e and f) performed in a similar manner to the bimodal case, where the second and third mode amplitudes of the recovered distributions were different to the test distribution. In both trimodal examples where the peak magnitudes are not equal, we observe the second distribution mode peak position has minor differences compared to the test distribution. Despite some overt variations between the recovered and test distributions, the key features (number of peaks, peak positions, peak width and peak magnitudes) are largely replicated and were well recovered using our regularised inverse method.

**Fig 2.**
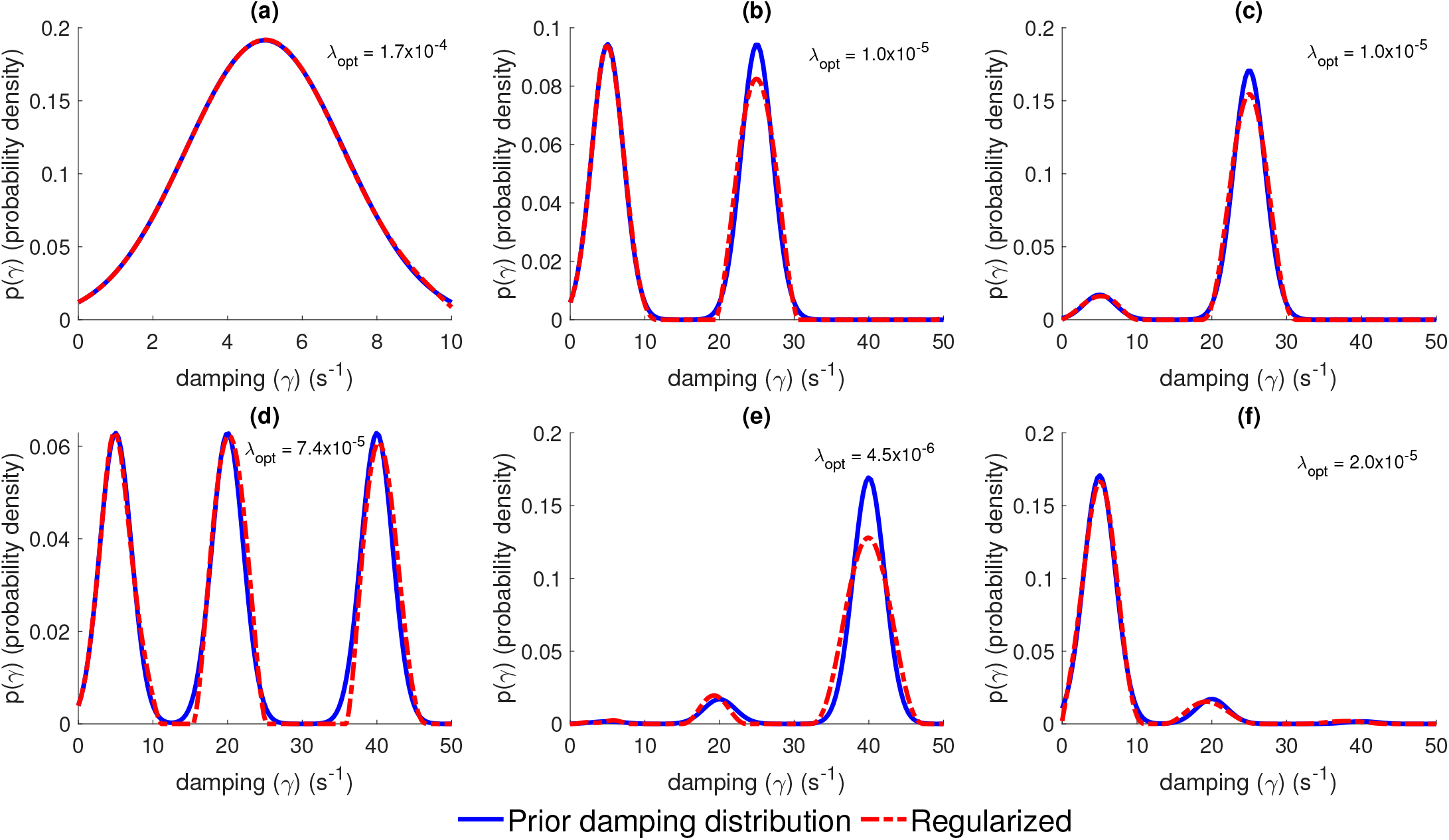
Validation of the inverse method by the recovery of prior damping distributions. **(a)** Unimodal prior damping distribution. **(b)** Bimodal prior damping distribution where both modes of same peak magnitude **(c)** Bimodal prior damping distribution where the relative magnitude of the peak of the most weakly damped mode is 0.1 that of the peak of the heavier damped mode. **(d)** Trimodal prior damping distribution where all modes are of the same peak magnitude. **(e)** As for (d) but relative peak mode heights are 0.01, 0.1 and 1 respectively. **(f)** As for (d) but relative peak mode heights are 1, 0.1 and 0.01 respectively. Blue lines are the prior distributions, whereas red dotted lines represent the recovered distributions using the inverse Tikhonov regularisation method and λ_opt_ are the optimally chosen regularisation parameters. For further details refer to Methods.

### Empirically estimated damping distributions

The empirically estimated damping distributions, across both EC and EO conditions were largely found to be bimodal, at 82% and 70% respectively. Examples of empirically recovered distributions are shown in Fig 3 along with the respective EC and EO condition power spectral densities. To determine whether the bimodal structure was a numerical artefact, we tested whether the removal of the second mode still permitted good spectral fits in the cases where multiple modes were present. The second distribution mode was removed by setting all relevant weightings to 0. The altered damping distributions were all renormalized to appropriately account for the removal of the second mode weightings and were then used in the forward model Eq 14 to generate spectral fits. We found that eliminating the second mode resulted in an inability to fit the high frequency scaling of the power spectrum, indicating that its presence was necessary to obtain optimal fits. As expected we found that power spectra with more pronounced alpha peaks were generally associated with damping distributions that were more weakly damped using the weakly damped measure described in Methods. The area of the un-normalized empirically estimated damping distributions were not significantly different between EC and EO conditions, suggesting that state induced spectral power changes are predominantly the results of changes in empirically estimated probability distribution shape (See S1 Fig for details). Further, this shape did not depend significantly on the exact form of the integral kernel used. Using the kernel of Eq (5) instead of Eq (6) made little difference to the shape of the empirically estimated damping distributions and no difference to the number of inferred modes (results not shown). It is entirely possible though, that using alternative kernels (Gaussian, Gamma) may result in distributions of differing structure, that may not present with the same bimodal distribution in dampings. However, these functional forms are not immediately compatible with the theoretical and experimental framework that we have used. As such, it would be more difficult to motivate their use over the approximate Lorentzian form employed here.

**Fig 3.**
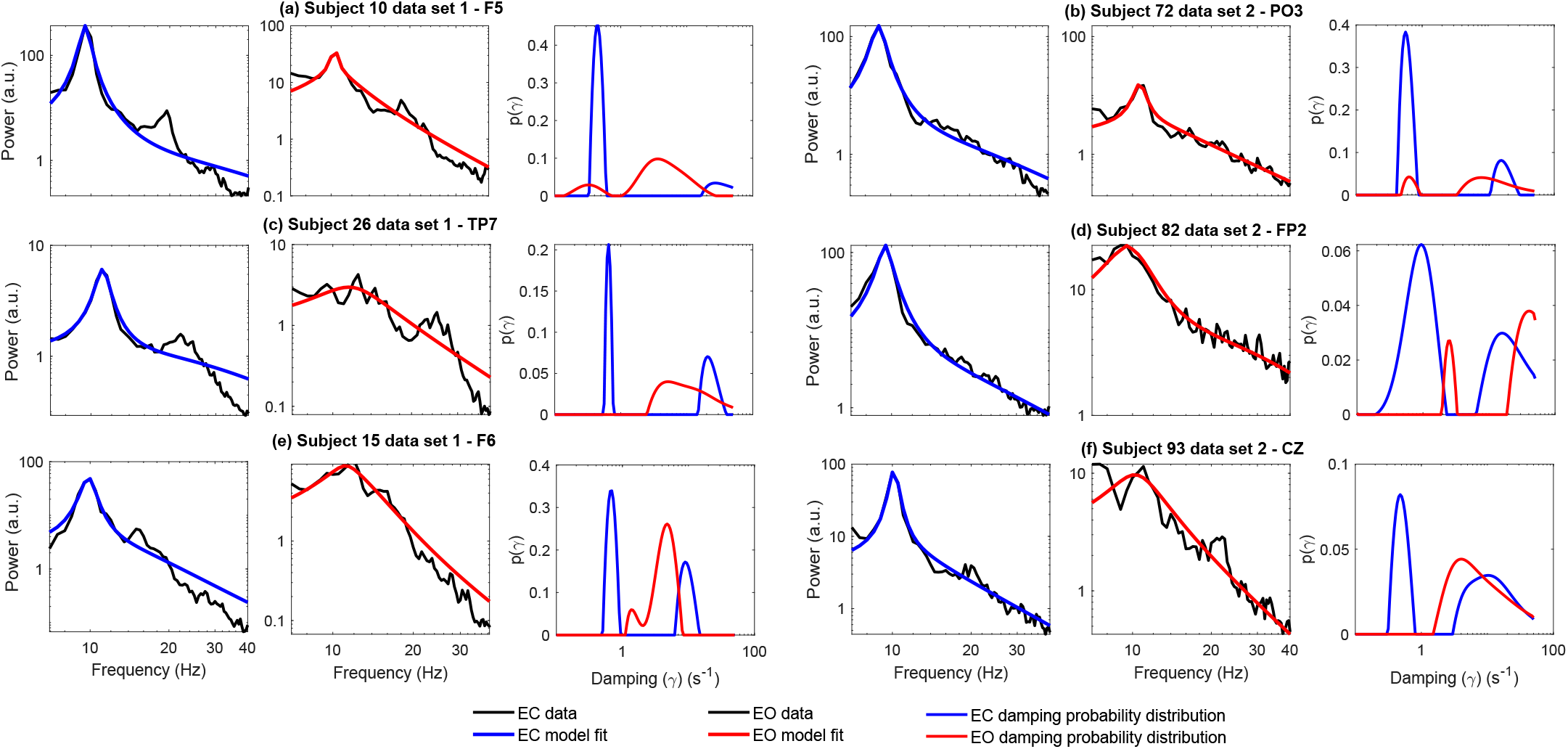
Example damping distributions for strong alpha blockers. **(a-f)** Damping distributions obtained from EC (blue) and EO (red) power spectra for six individuals (three per data set) who presented with strong alpha blocking. Plotted alongside the experimental power spectra are the respective model fits (EC - blue, EO - red) generated using the EC and EO damping distributions in the forward problem Eq 14. The inferred damping distributions are clearly bimodal in shape, a feature common across both data sets. The first mode of the EC damping distribution is generally peaked at a smaller damping value and has a larger respective area when compared to the EO distributions.

### Spectral fit

By using the empirically estimated damping distributions in the forward problem (i.e Eq 14) to generate model spectra, we are able to demonstrate that we are able to obtain good spectral fits that replicate the power-law like scaling measured in the original experimental data. All the model spectra were generated using the maximum entropy damping distributions obtained by evaluating a range of regularisation parameters with fit quality evaluated using a RSS between the original EEG spectra and the forward model spectra. Fig 4a presents the resulting RSS distribution of model deviation for both data sets in a pooled analysis. Spectral plots in Figs 4b illustrate examples of a range of fit quality. We analysed the distribution of RSS to gain insight into the fit quality most prevalent in our model. The median of the distribution (EC and EO) corresponds to model fits that are of high quality. Even in instances of poor fit qualities (bottom 5% of fit quality) the experimental spectra are reproduced reasonably well. In general the shape of (in particular alpha peak location, amplitude and high frequency scaling) most power spectral densities could be well accounted for as a sum of damped alpha band oscillatory processes. However the presence of significant resonant beta band (13 - 30 Hz), which was not explicitly accounted for in our approach, was a major contributing factor to poor model fit. We discuss this issue in more detail in the Discussion.

**Fig 4.**
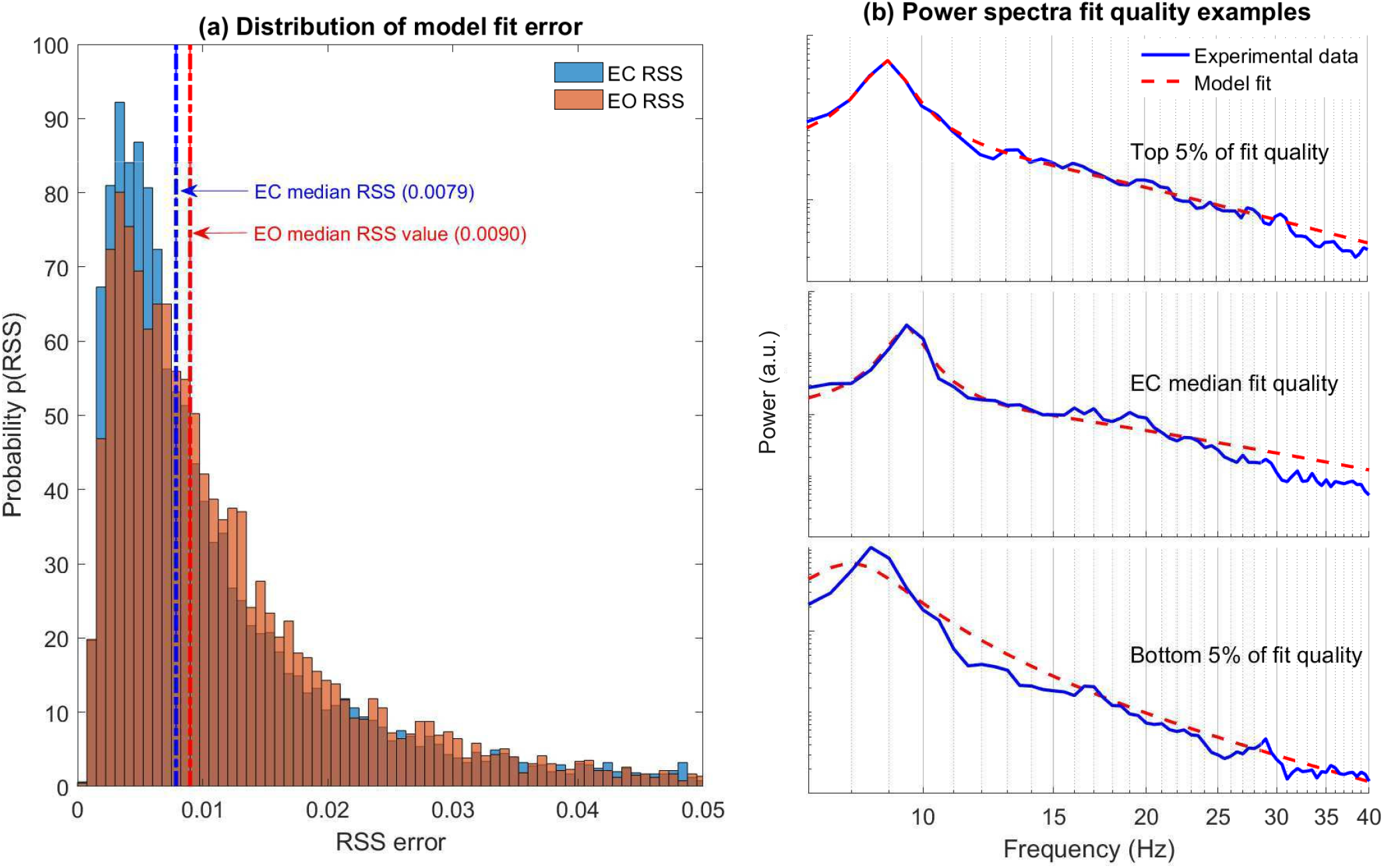
Resting EEG power spectral densities are well described by a sum of damped alpha oscillatory processes. **(a)** Histograms of the minimum residual sum of squares (RSS) error between experimental and model spectra for EC and EO states in pooled data set analysis. The median *RSS* error of *EC* = 0.0079 and *EO* = 0.0090 are associated with objectively good fits. **(b)** Examples of model fit quality for three distinct cases. Plots are ordered top to bottom with decreasing fit error (EC spectra presented).

The power spectrum scaling exponents, *β*, were computed across all subjects and channels for both the experimental spectra and the model by fitting a 1/*f^β^* profile over the frequency range 15-40 Hz. Pooled data (*N* = 8677 *β* values for experimental and model spectra across each state) analysis revealed a range of experimental spectral scaling across EC and EO conditions (Fig 5a) with the most probable values of *β_E_* ≈ 1.8 and *β_E_* ≈ 1.7 respectively. The broad range of scaling behaviour recorded in the experimental power spectra was reproduced in the forward model spectral fits (Fig 5b). Across the pooled data the model and experimental spectral exponents are well correlated in EC (*r* = 0.64, *p* = 10^−6^) and EO conditions (*r* = 0.81, *p* = 10^−6^) (Fig 5c).

**Fig 5.**
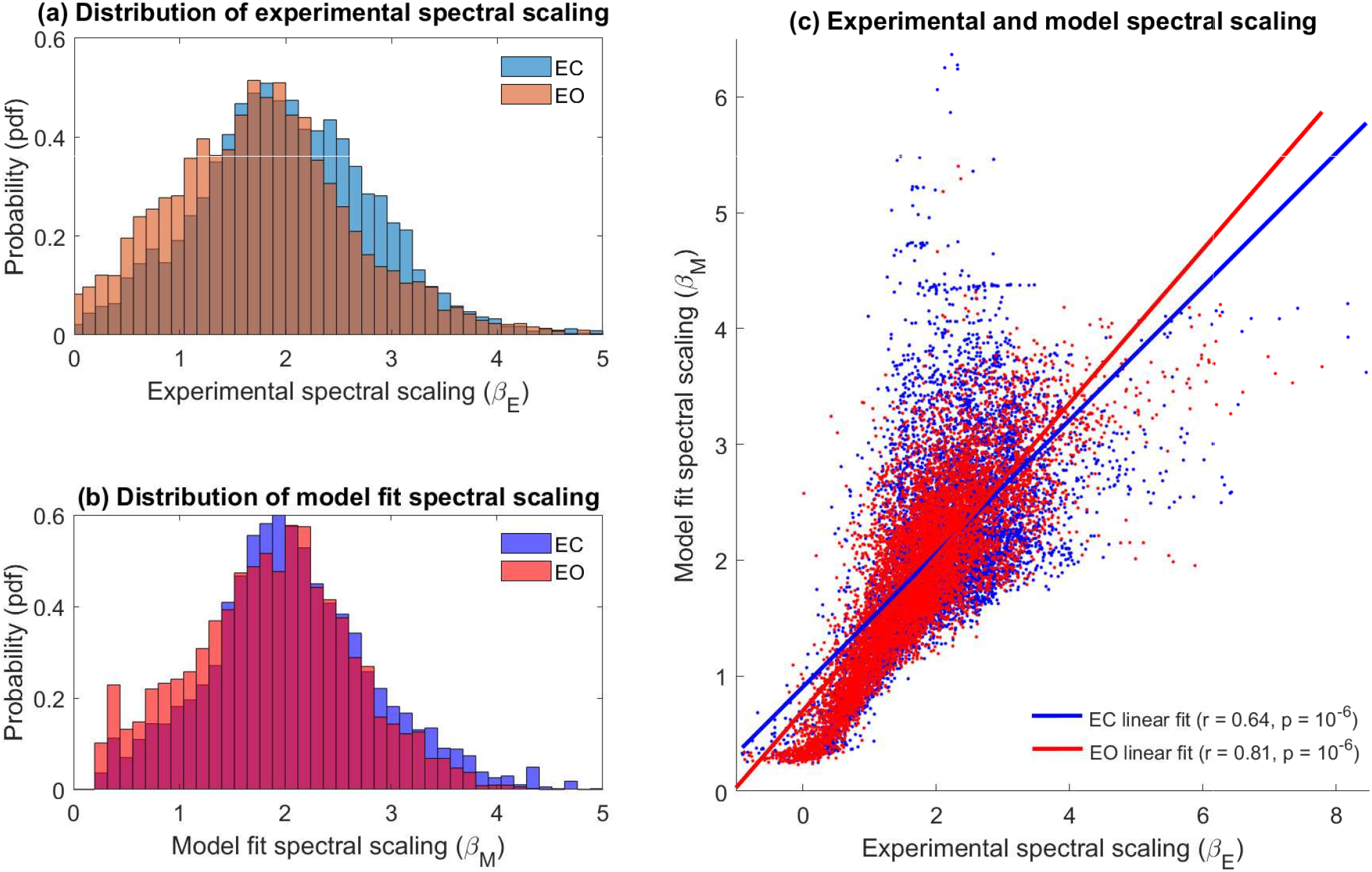
Experimental and model spectral scaling in EC and EO states. **(a)** Probability distribution of experimental spectral scaling exponent (*β_E_*) for EC (blue) and EO (red) states with data set 1 and 2 pooled for analysis. **(b)** Same as (a) but for optimal model fit spectral scaling exponent (*β_M_*). **(c)** Scatter plot of the experimentally determined spectral scaling parameter (*β_E_*) versus the model calculated spectral scaling parameter (*β_M_*) for EC (dots - blue) and EO resting state (dots - red). The experimental and model fit scaling parameters are well correlated across both recording conditions (EC: *r* = 0.64, *p* = 10^−6^ / EO: *r* = 0.81, *p* = 10^−6^). Statistical significance calculated via a nonparametric permutation test.

### Alpha blocking in resting state EEG

To quantify alpha blocking we used resting state data recorded in EC and EO conditions from two separate EEG data sets having a combined total of 136 participants, and computed channel based power spectral densities using Welch’s method. A total of 8677 (109 subjects × 64 channels + 27 subjects × 63 channels) power spectra were estimated for each recording condition. The Jensen-Shannon divergence was then computed between each EC/EO spectral pair to measure the degree of alpha blocking across the respective data sets. A range of alpha blocking was observed across the both data sets; some participants evinced little blocking whereas others exhibited strong alpha band attenuation between states. Example cases are plotted in Fig 6a-f where power spectra from an occipital electrode (O1) are plotted together in descending order of the degree of alpha blocking. Fig 6g-h shows the topography for two strong blockers exhibiting classical occipital dominance. Such an occipital dominance was clearly seen in the respective group averages (Fig 6i-j).

**Fig 6.**
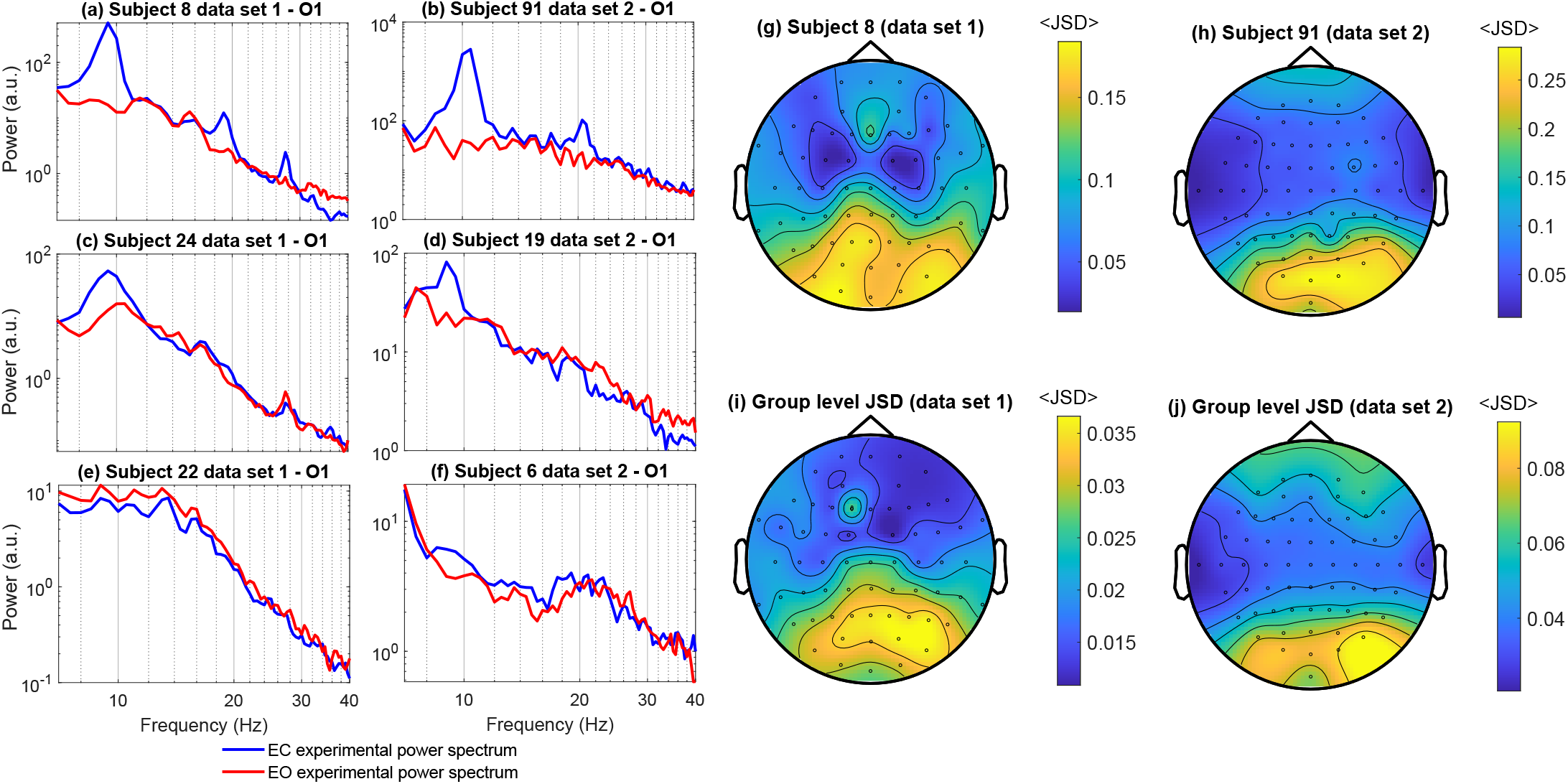
Spectral and topographic properties of alpha blocking in healthy participants. **(a-f)** Examples of spectral variation between EC and EO resting states (O1 electrode) with three subjects shown for each data set. Plots are ordered top to bottom with decreasing alpha blocking (JSD). The exemplar cases demonstrate the contrast between strong and weak alpha blockers. **(g-j)** Topographic maps showing the occipital dominance of alpha blocking for a typical individual (g-h) and at the group level (i-j) across data set 1 and 2. Group level average is computed across subjects on an electrode-wise basis. Note, in order to highlight the consistent pattern of occipital dominance topographic maps are all displayed using different scales.

### Changes in damping distributions can account for alpha blocking

Our estimated damping distributions suggest that alterations in the alpha peak power (see Fig 3) were predominantly driven by changes in the weakly damped mode. We therefor quantified this relation further by investigating the spatial distribution of both the JSD and the group-level differences in damping. Changes in the damping distribution between EC and EO conditions were quantified by calculating the difference in weakly damped measure (Eq 25) between EC and EO states (i.e. *WDM_EC_* – *WDM_EO_*), on an electrode by electrode basis for both data sets. Group level averages were computed and plotted topographically such that direct comparisons can be made between the topography of alpha blocking quantified using the Jensen-Shannon Divergence (see Fig 6i-j).

As expected group-level weakly damped measure difference values were largest in the occipital regions for both data sets (Fig 7a and b). However, in addition to a clear occipital dominance, statistically significant differences in the weakly damped measure were topographically widespread, particularly in the second data set. The similarity of the topographic variation of the difference in the weakly damped measure between EC and EO states and the Jensen-Shannon Divergence (Fig 6i-j) are clearly apparent. Averaging across subjects in each data set and plotting electrode-wise we see (Fig 7c) that these respective measures are well correlated across data set 1 (*r* = 0.838, *p* = 10^−6^) and data set 2 (*r* = 0.901, *p* = 10^−6^) On this basis we can reasonably conclude that alpha blocking is being driven by increases in the damping of a weakly damped population of alpha band oscillatory processes.

**Fig 7.**
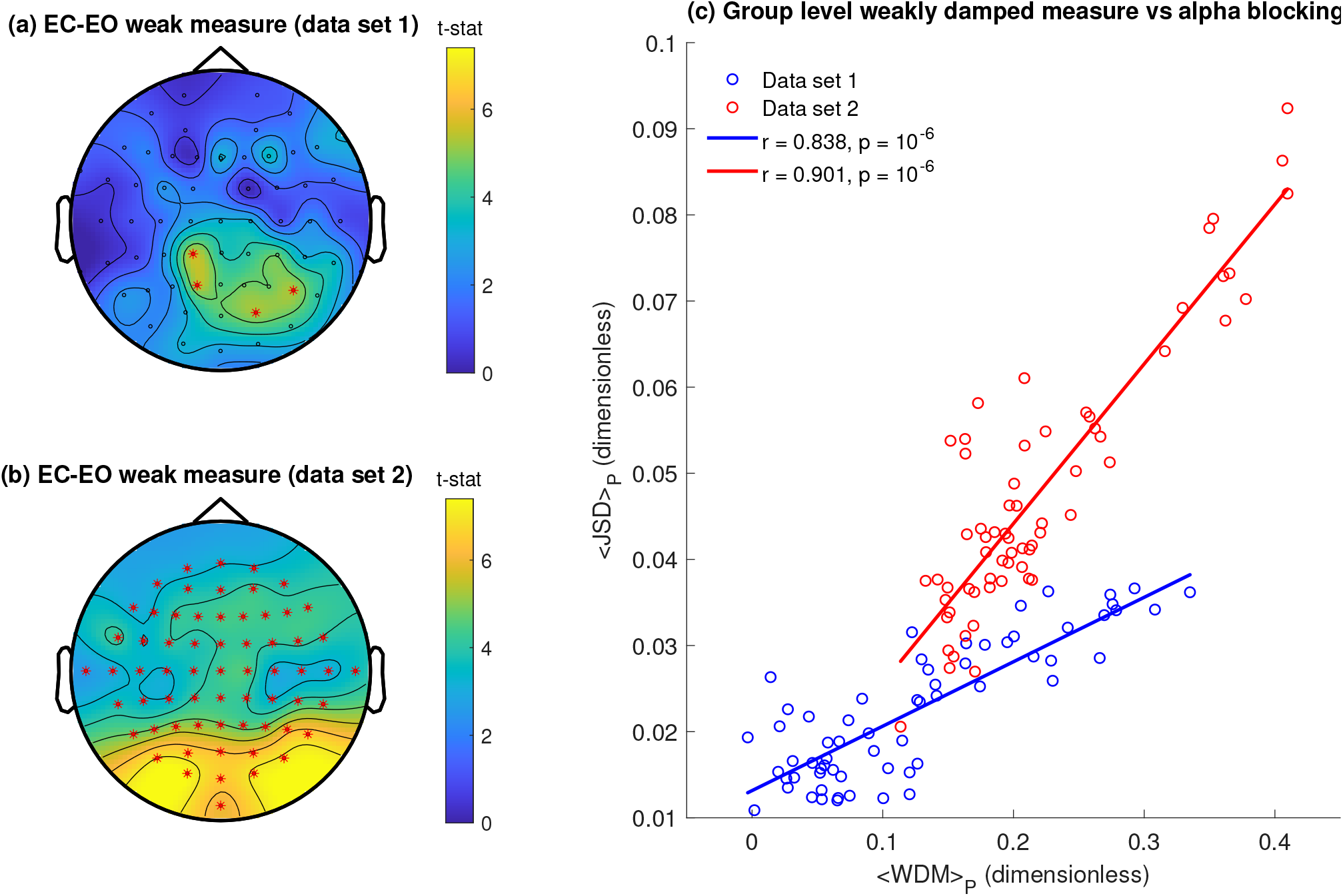
Alpha blocking is driven by changes in the distribution of alpha oscillatory damping. Topographic maps of mean difference between the weakly damped measure in EC and EO states **(a)** Data set 1 **(b)** Data set 2. Red asterisks indicate significant (*p* < 0.05) differences in the weakly damped measure between EC and EO states calculated using the Max/Min-t permutation test (see Methods) and corrected for multiple comparisons. **(c)** Scatter plot of electrode-wise Jensen-Shannon divergence and the difference in the weakly damped measure between EC and EO, averaged across all participants in each data set (red = data set 1, blue = data set 2). In general the two measures are well correlated with each other: data set 1, *r* = 0.838, *p* = 10^−6^; data set 2, *r* = 0.901, *p* = 10^−6^. Note that the group-level quantities differ between the data sets, in particular the <JSD>. The reason for this is not clear. It is possible that it is due to differing experimental configurations of the two studies, or more likely, it is due to a larger prevalence of strong alpha blockers in data set 2 (Fig 6j). Statistical significance was validated using a nonparametric permutation testing.

## Discussion

Our aim was to explore whether the resting EEG could be accounted for in terms of a sum of damped alpha oscillatory processes and in particular whether changes in the power spectrum between EC and EO states could be reasonably accounted for solely by changes in the damping of such oscillatory processes. Through the use of regularisation methods, we estimated EC and EO damping distributions across a total of 136 participants, available from two independently collected EEG data sets, and characterised their features. We found the model performed well when applied to real EEG spectra, providing high quality fits to both the alpha band and the high frequency power-law-like scaling. The attenuation of the peak alpha power observed in heavy alpha blockers seems to be well explained by an increase in damping between EC and EO states as we found that the distribution of damping became more heavily damped in EO states than in EC when alpha blocking was present. Our results therefore suggest an economical approach to describing the resting EEG spectrum in terms of the summed activity of alpha band damped linear oscillations and represents an alternative way to characterising the spectral properties of resting EEG.

Our findings suggest both alpha blocking and ‘1/*f*’ noise in resting EEG, in contrast to the dominant view, might be accounted in terms of a single underlying mechanism. This is significant as the mechanistic genesis for the alpha rhythm and its EC-EO induced remains uncertain. The received view for the origin of the alpha rhythm is that it arises principally via reverberant thalmo-cortical neuronal population interactions [7].

In contrast the predominant explanation for alterations in alpha band power, in response to various tasks and changes in behavioural state, are changes in the dynamical synchronization of microscopic neural population activity [14]. Using a model that is theoretically founded in mean-field model approaches, which assumes the EEG signal is generated by the aggregate activity of multiple coupled excitatory and inhibitory cortical neural populations, we have shown that it is possible to account for alpha blocking in terms of changes in the underlying damping of stochastically driven alpha band damped linear oscillations. This approach inherently incorporates a description of ‘1/*f*’ noise in EEG by including a distribution in the dampings of the alpha oscillatory processes. On this basis viewing the resting EEG as being composed of rhythmic processes occurring on a background of arrhythmic activity, suggested to be scale-free in nature and indicative of self-organised criticality [25], is not necessarily required. The heterogeneity of spectral scaling seen in resting state EEG (see Fig 5) can be readily accounted for without the need to invoke the presence of fractal temporal dynamics or other non-linear processes. We have provided empirical evidence showing that it is possible to account for both alpha blocking and ‘1/*f*’ noise through a single underlying mechanism.

The most surprising result we obtained was the predominant bimodal structure in the distribution of dampings. Interpreting this from a model fit perspective though is in hindsight fairly simple i.e. two modes are required to achieve a superior fit to both the alpha peak and the ‘1/*f*’ tail. From a neurophysiological perspective however, interpretation of these two persistent distribution modes is more challenging. One interpretation would be that large changes in damping could be indicative of a more responsive neural system. We found that in instances where a large degree of alpha blocking was present, the mode separation was largest in the EC state which was subsequently reduced in EO. In some cases, distributions went from bimodal to unimodal in EC and EO states for particularly strong alpha blockers. This could mean that participants who are subject to a large difference in damping between states, are more efficient at corralling the underlying neural population activity into task engagement and then subsequently dispersing it when no longer needed. When the eyes are closed the brain is in a more restful idling state where a larger proportion of the cortical neural populations are weakly damped. Upon opening of the eyes, most neural population activity is suppressed in response to incoming stimuli while a small undamped proportion is engaged in the task, as reflected in systematic large scale increases in the damping. Closing the eyes once more engages neuronal population activity and sets them to idling as the demand for cognitive resources reduces. Such a view lends support to the notion that highly reactive alpha band activity is correlated with better memory engagement and learning/intelligence [42, 43, 43, 44]. Indeed, research has shown that alpha power and peak frequency respond in a variety of ways to mental effort and memory retention and/or retrieval [11]. While this interpretation is highly speculative, it does provide a clear motivation for further explorations of the functional significance of our approach. For example we could assume that our damping distribution can be well represented as a sum (or mixture) of one or more parametric distributions with our goal being the optimal estimation of the corresponding parameters. In contrast we assumed no prior functional form for our damping distributions.

### Model limitations and extensions

While the simplicity of our model is one of the features that makes it useful, there exists some clear limitations that were not addressed within this work, those being the low frequency scaling of the power spectrum and presence of clear resonant beta band (13 - 30 Hz) activity. Resting EEG has often been characterised as exhibiting two frequency scaling regions, one in the low frequency and one high frequency with the transitional region occurring somewhere in the alpha band [18, 28]. For the sake of simplicity, we neglected the low frequency scaling and chose to investigate a broadband region starting in the alpha band. Future work needs to extend our model to include either a single low frequency relaxation oscillation or a distribution of such processes. This inclusion would then posit that the resting EEG is almost entirely explained by the joint activity of low frequency and alpha band processes that have a distribution of relaxation rates. In support of this idea Chaudhuri et. al. (2018) [39] demonstrated that broadband ECoG power spectrum could be modeled by the sum of two Lorentzian processes, suggesting that the underlying neural activity had two distinct time courses with dominant low frequency and high frequency components.

Our exclusion of beta band activity could be rectified in two ways. The simplest would be to simply filter out the beta band activity and attempt to maintain the scaling present in the spectrum. This would correct any errors introduced to the model fit when attempting to model the spectrum when a large beta peak is present. However, beta band activity is typically highly correlated with the alpha band, with some hypothesizing an explicit harmonic relationship [45, 46]. On this basis it would be better to modify the underlying model assumptions to incorporate beta band activity. However, it is not immediately clear how this would be achieved and any such alteration would undoubtedly complicate the model by suggesting, possibly non-linear, interactions between damped alpha and beta band processes.

One final issue is that we assumed that EC-EO changes are solely driven by changes in damping. However changes in the peak alpha frequency between EC and EO states are often observed [11]. In the current study the peak alpha frequency was selected by peak fitting each condition (see Methods), thus the change was incorporated before the model was applied to the fit. A general solution would be to extend the model to include a distribution of frequencies in addition to dampings, resulting in a two dimensional probability density function over both alpha frequency and damping. However, such inclusions complicate the problem substantially. Nevertheless, preliminary analyses, not reported here, indicate that it may be possible to solve such a problem using gradient descent methods.

We have demonstrated that alpha blocking and ‘1/*f*’ noise in resting EEG can be accounted for by a simple singular mechanism consisting of damped alpha band oscillatory activity. Through the use of inverse methods, the distribution of damping was typically revealed to be bimodal in both EC and EO resting states, with the degree of alpha blocking measured between EC and EO shown to be driven by systematic increases in the damping of a weakly damped mode. The topographic distribution of these changes in damping paralleled well the corresponding topographic changes in alpha blocking. The power-law scaling of the generated spectral fits were shown to be well correlated with the corresponding experimental power spectra being able to reproduce the 1/*f^β^* tail. While our assumption that the EEG signal is composed of the bulk collective activity of many uncorrelated stochastic alpha band damped linear oscillatory processes is simplistic it does provide a novel way of interpreting the spectral behaviour of resting EEG.

## Methods

### EEG as a sum of damped relaxation oscillatory processes

The fundamental assumption of our model is that the resting EEG signal (measured at the sensor level) is generated by a vast number of uncorrelated stochastically perturbed alpha band damped linear oscillations that arise from multiple neural populations, which have a distribution of dampings (relaxation rates), the form of which is unknown to us. Our assumed model is theoretically motivated by a general model for generating 1/*f* noise [16] and by mean-field approaches to modelling the resting EEG [29].

We begin by noting that most linearised mean-field models generally account for the EEG signal, eeg(*x,t*) in a quantitatively similar manner i.e. a linear time invariant transfer function driven by broad-band noise

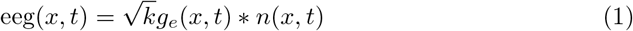

where *g_e_*(*x,t*) is the (*excitatory*) *electrocortical impulse response* [33,34] for cortical location *x, n*(*x, t*) is the driving broad-band noise process, * is the convolution operator and 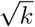 is a constant taking into account the (bio)physical processes associated with EEG recordings. In the frequency domain this becomes

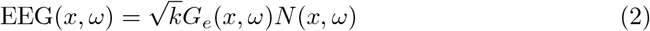

where the conjugate variables are EEG(*x,ω*), *G_e_*(*x,ω*) and *N*(*x,ω*). Stable linearisation requires an impulse response that has a decaying envelope. Therefore, we choose the simplest parametric electrocortical impulse response function as a damped cosinusoid assuming it has an instantaneous rise time in response to perturbation (rise time is much shorter than the characteristic decay rate). On this basis the functional form is

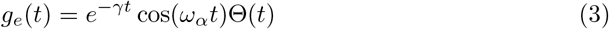

where the spatial dependency on *x* has been removed for clarity, *ω_α_* = 2*πf_α_* is the oscillating frequency with *f_α_* the (parametric) centre alpha frequency, *γ* is the parametric damping and Θ(*t*) is the Heaviside step function (it is entirely expected that *γ* and *f_α_* will depend on a set of population model biophysical parameters [*γ* ≡ *γ*(*p*), *f_α_* ≡ *f_α_*(*p*)], however the current parameterisation is ignorant to this). From Eq (2), the power spectral density is then

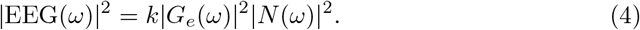

By assuming the electrocortical impulse response of Eq (3), |*G_e_*(*ω*)|^2^ can be calculated as:

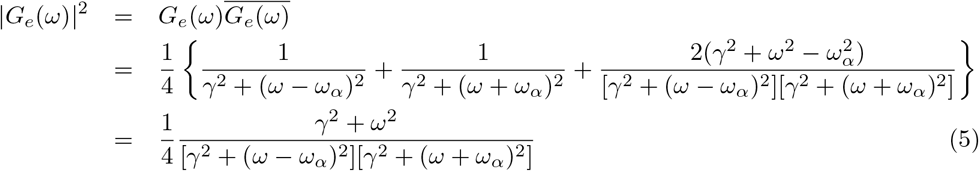

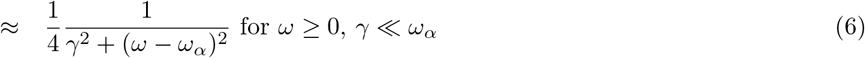

where 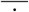 is the complex conjugate. Rewriting in terms of *f_α_*, Eq (6) (given that there is uncertainty about the exact form of *g_e_*(*t*) and our choice is merely the simplest, we choose the numerically simpler reduction of Eq (6) instead of Eq (5)) is then

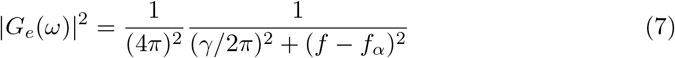

We now assume that the signal measured at a single location is the sum of many such processes arising from multiple neural populations, where the centre alpha frequency, *f_α_* is assumed constant and remains physiologically fixed, while the damping *γ* varies:

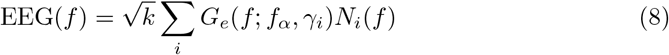

The estimated power spectral density for this model process, *S*(*f*), is then,

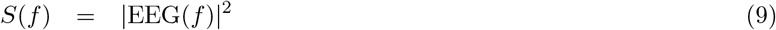

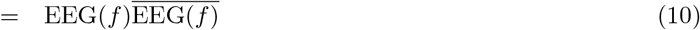

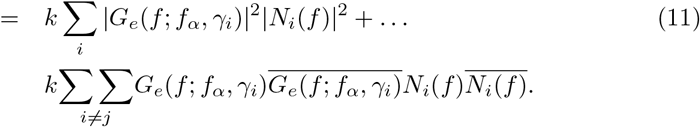

We now assume

- the white noise driving processes *N_i_*(*f*) are all uncorrelated such that their cross-spectrum vanishes,
- the *N_i_*(*f*) are broadband and of equal power (RMS) i.e. *N_i_*(*f*) → *N* (i.e. is a constant),
reducing our estimated model power spectral density to

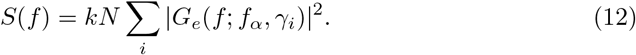

We now make one final assumption that the total number of damped linear oscillators, *M*, is sufficiently large that we can define a distribution of dampings, *p*(*γ*), such that the number of oscillators with damping between *γ* and *γ* + *dγ*, is *Mp*(*γ*)*dγ*, where ∫*p*(*γ*)*dγ* = 1. On this basis Eq (12) can be rewritten in the continuum limit as

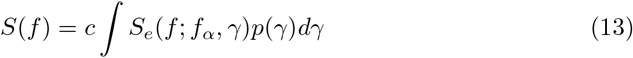

where *c* = *kNM* and *S_e_*(*f; f_α_, γ*) ≡ |*G_e_*(*f; f_α_, γ_i_*)|^2^. For our purposes *S*(*f*) will be empirically estimated directly from EEG data and *S_e_*(*f; f_α_, γ*) is our theoretically specified spectral response (Eq 7). Substituting this into our equation we arrive at our final model description

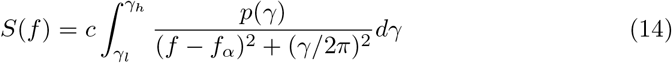

which is a Fredholm integral equation of the first kind [35]. In practice, the peak alpha frequency is selected as described in the Methods section and the upper (*γ_h_*) and lower (*γ_l_*) limits of the damping distribution are chosen such that a broad range of spectral behaviour is covered i.e. both large and sharp alpha peaks and those which are heavily damped.

### Numerical solution of the model Fredholm Integral equation using Tikhonov regularization

Fredholm integral equations of the first kind are inherently ill-posed and difficult to solve [36]. For this reason to obtain useful results, regularization methods are required. We employ the use of Tikhonov regularization, a well known and studied method for solving ill-posed problems [36]. The process used to estimate the damping distributions using Tikhonov regularization is described as follows. Beginning with our model formulation, Eq (14), we define our Lorentzian-shaped frequency domain electrocortical transfer function as

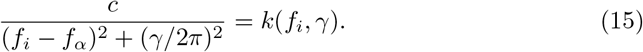

Our general Fredholm integral equation of the first kind is then

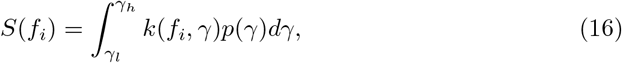

where we note that we have incorporated our constant c into Eq (15). Making one further simplification by setting *k_i_*(*γ*) = *k*(*f_i_, γ*), we then discretize the integral

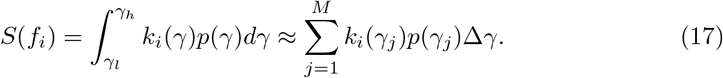

Finally, we can arrange the above summation as a set of linear equations in matrix notation

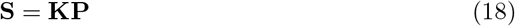

where **S** is an *N* × 1 data vector (power spectrum), **K** is an *N* × *M* coefficient matrix with entries consisting of the kernel evaluated at each damping value and **P** is the unknown distribution vector *M* × 1 in dimensions. Simply inverting the equation

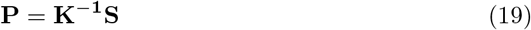

leads to numerically unstable solutions that are sensitive to variations in **S** and those caused by the numerical inversion. By applying appropriate regularization to Eq (18), we can obtain sensible solutions from solving the inverted equation. Solving the Tikhonov regularization problem involves finding appropriate values for **P** that minimizes the following function for

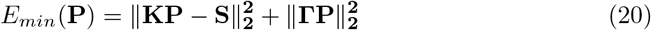

where Γ = λ**I** is the *Tikhonov Matrix*, which is generally chosen to be identity matrix, **I**, multiplied by a regularization parameter, λ and ||·|| is the Euclidean (or *L*^2^) norm. In practice, we solve Eq (20) by setting it up as a constrained general least squares minimization problem of the form

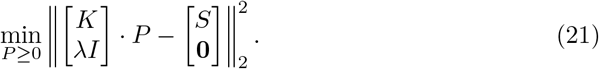

and use MATLAB’s lsqnonneg algorithm. With all problems requiring regularization the difficulty in obtaining an appropriate solution is further compounded by the determining the ‘optimal’ regularization parameter. The degree to which a solution is regularized is ultimately a trade off between over or underfitting. When the magnitude of the regularization parameter is large, the determined solutions will be overly smooth. When the regularization is small, the solution will still be noisy and susceptible to the effects of numerical noise. Methods exists that allow an optimal regularization parameter to be determined, but these are largely heuristic with optimal defined as the value which satisfies the constraints of the experiment in question [36]. For the current study, we explore a regularization parameter range of 10^−5^ ≤ λ ≤ 10^0^ divided into 100 logarithmically spaced steps, with the optimal regularization parameter chosen such that it finds the maximum entropy solution that falls within an error bound of approximately 2.5% of the minimum residual sum of squares error (RRS) between the generated power spectrum and the data power spectrum. The maximum entropy distribution within that error bound was chosen as it favours distributions that are smoother and broader. See S2 Appendix for a graphical example of the optimal regularization parameter selection process.

### Simulated tests

A simulation study was conducted to validate the Tikhonov regularisation method. Damping distributions of known form that were uni/bi/tri-modal with differing relative mode areas were used in the forward problem to generate simulated power spectra. The simulated power spectra were then used as the input to the regularised inversion method of the previous section to recover the known distributions. The simulated damping distributions were Gaussian in form

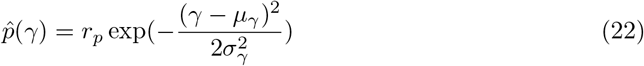

with *r_p_* the ratio between the peak magnitudes, *μ_γ_* the mean and *σ_γ_* the standard deviation. For bi/trimodal structures, the damping distributions were produced by summing multiple Gaussians. The relevant model parameters tested are presented below:

- **Unimodal:** Damping interval [0 ≤ *γ* ≤ 10] s^−1^, *μ_p_* = 5 s^−1^, *σ* = 5 s^−1^.
- **Bimodal:** Damping interval [0 ≤ *γ* ≤ 50] s^−1^, *μ*_1_ =5 s^−1^, *μ*_2_ = 25 s^−1^, standard deviation *σ* = 5 s^−1^ and *r_p_* = (1, 1) and (0.1, 1).
- **Trimodal:** Damping interval [0 ≤ *γ* ≤ 50] s^−1^, *μ*_1_ = 5 s^−1^, *μ*_2_ = 20s^−1^, *μ*_3_ = 40 s^−1^, *σ* = 5 s^−1^ and *r_p_* = (1,1,1), (0.01, 0.1, 1), (1, 0.1, 0.01).

A successful recovery of a known distribution was measured using a RSS cost function between the known distribution and the recovered distribution. A regularisation parameter range of [10^−^6 ≤ λ ≤ 1] was used and divided into 100 logarithmically spaced steps.

### EEG data

Two freely available EEG data sets collected from two independent experiments were chosen because both explicitly included EC and EO resting condition recordings. The first set (subsequently referred to as Dataset 1) was made available by [37] and consisted of data from 30 participants recorded across multiple drug/task conditions during EC and EO resting state across 64 channels using a modified 10-20 layout. We used the data of 27 participants in the EC and EO pre-drug conditions. Each EEG recording was approximately 2 minutes in duration and was recorded at 500 Hz. Three participant recordings were not used due to significant artefact. The second set (subsequently referred to as Dataset 2) is an open source EEG data made available by [38] (https://physionet.org/content/eegmmidb/) and includes 109 participants who had their EEG recorded, using the BCI2000 system, across 14 experimental tasks. We make use of the 2 minute EC and EO recordings that were collected from 64 electrodes using a modified 10-20 layout at a sampling frequency of 160 Hz. We chose not to group data sets together for analysis as each dataset used slightly different channel layouts.

### Power spectral analysis

All spectral analysis was performed at the sensor level. Power spectra for the EEG was calculated by computing the Welch periodogram across all channels using the measured time series with 2 second Hamming windows with 50% overlap. Power spectra broadband activity between *γ* Hz and 40 Hz is used for the EEG data as it encompasses alpha band activity and the high frequency scaling which are our main focus. The low frequency cut off was chosen as there is theoretical and empirical evidence to suggest that the low frequency scaling of the power spectrum is generated by a low frequency resonance which we do not model here [34, 39]. Thus we choose to exclude it from the spectral interval of interest. The upper frequency cut-off was selected to avoid 50/60 Hz mains line activity and to avoid EEG spectra that tend towards a white noise profile at higher frequencies. The peak alpha frequency, *f_α_*, for each experimental spectrum was calculated by fitting a single function of the form 1/[(*f* – *f_α_*)^2^ + *b*^2^] within the alpha band using MATLAB curvefit toolbox functions. Because a comparison between the experimental and model spectral slope for frequencies is explored, a spectral scaling parameter *β* was calculated by fitting a 1 /*f^β^* profile to the power spectrum over a frequency interval from 15-40 Hz. This procedure was performed on all the experimental spectra and their respective model spectra.

### Quantifying alpha blocking using the Jensen-Shannon Divergence

The degree of alpha blocking present between EC and EO conditions varies between participants: some presenting with significant alpha attenuation and others showing minimal blocking. To explore and characterize the presence of alpha blocking within the EEG data we require a measure that can quantify the degree to which an individual attenuates alpha band activity between EC and EO states. On this basis we employ the same methodology used by [32], where the Jensen-Shannon divergence (JSD), *D_JS_*, between the respective EC and EO power spectra is used to quantify the level of alpha blocking. The JSD is a symmetric and non-negative measure of the distance between two probability distributions. By normalizing the power spectra to unity, we can treat each as a probability distribution allowing us to compute the distance between the EC and EO spectra in probability space. The JSD provides a useful way to measure the degree of alpha blocking between resting states. The expectation is that the larger the attenuation of alpha band activity between EC and EO states, the larger the JSD will be between the two distributions. The JSD is based upon the Kullback-Liebler divergence [40], *D_KL_*, which is defined as follows;

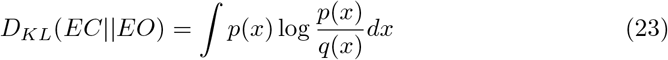

where *p*(*x*) and *q*(*x*) are two probability distributions. The JSD for two spectra is then

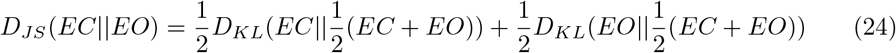

### Weakly damped distribution measure

To explore changes in the damping distributions between EC and EO states requires a measure that can quantify how ‘weakly damped’ a given distribution is. An obvious choice would be the peak position of the distribution. However, in instances when multiple distribution peaks are present, this simple method does not accurately characterise the dominant decay rate. Furthermore, given that the respective weighting of each damping plays an essential role in the outcome of the spectral profile, the peak position alone is not sufficient to quantify which distribution is more weakly damped when an overlap in the distribution peak may occur across conditions. Therefore, we constructed a measure that takes into account both the peak position and the respective area that lies under each mode in each recording condition with respect to the other state. The measure can be computed for any given pair of distributions and assigns a specific value to each resting state. A distribution (i.e *p*(*γ*) in Eq 13) is considered more weakly damped if it contains a larger area that falls to the left side of the other distributions peak. This measure is applied to the distribution peak or first mode if the distribution is multimodal. In practice, calculating the weakly damped measures (WDM) for two distributions requires calculating two specific values for each resting state damping distribution, achieved as follows;

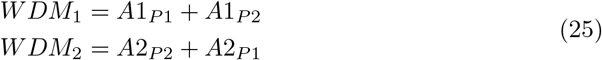

where *A*1_*P*1_ is the area in distribution 1 that falls before the peak of distribution 1, *A*1_*P*2_ is the area of distribution 1 that falls before the peak of distribution 2, with *A*2_*P*2_ and *A*2_*P*1_ being similarly defined. The weakly damped measures are then the sum of the two quantities for each state. The two values can then be compared and that which is larger is deemed more weakly damped i.e. *WDM*_1_ > *WDM*_2_ implies distribution 1 is more weakly damped than distribution 2. The inclusion of *A*1_*P*1_ and *A*2_*P*2_ terms are to account for situations comparing bimodal distributions where the first mode in one condition is much smaller than in the other (specifically with respect to weighting/area of the first mode) but happens to be positioned to the left of the other distributions first mode peak. In these instances, without the inclusion of these additional terms, the distribution with the smaller mode would have a weakly damped measure that was non-zero and the other distribution, which may have a much larger first distribution mode, would have a measure that is equal to 0, due to the way the *A*1_*P*2_ and *A*1_*P*2_ are defined. A graphical example of this and a further discussion is provided in S3 Appendix.

### Characterisation of numerically calculated damping distributions

For numerically estimated damping distributions the following features were quantified:

- Number of modes – the number of peaks, calculated using MATLABs peakfind function, in the respective EC and EO damping distributions.
- Mode positions – the peak position of each mode.
- Difference in damping between EC and EO states – calculated subject and electrode-wise as the difference between the respective weakly damped measures.

These quantities were calculated for both resting states across all subjects and channels providing a total of 8677 damping distributions [(109 Subs × 64 Channels) + (27 Subs × 63 Channels)] for EO and EC conditions. Given the differing channel layouts of the respective data sets separate group level topographic plots were calculated with any summary statistics being computed separately across the data sets.

### Permutation testing

We employed permutation methods to test whether there exists statistically significant differences between EC and EO resting state data. We use EEG channel based data averaged across participants and compute t-values from the difference in means between the two states. For each EEG sensor a null hypothesis, *H*_0,…,*i*_ is assumed in which there is no significant difference between EC and EO resting states. Given that multiple hypothesis tests are being conducted in parallel, the permutation test was corrected for the multiple comparison problem. Here we use the Max-t/Min-t method detailed in [41] which corrects for the inflated Type I error rate introduced by multiple comparisons. For our experimental permutation testing we permuted the data 5000 times. For the sake of clarity we provide a brief description of the Max-t/Min-t algorithm:

#### Algorithm 1: Max-t/Min-t Multiple Comparison Correction Permutation Test

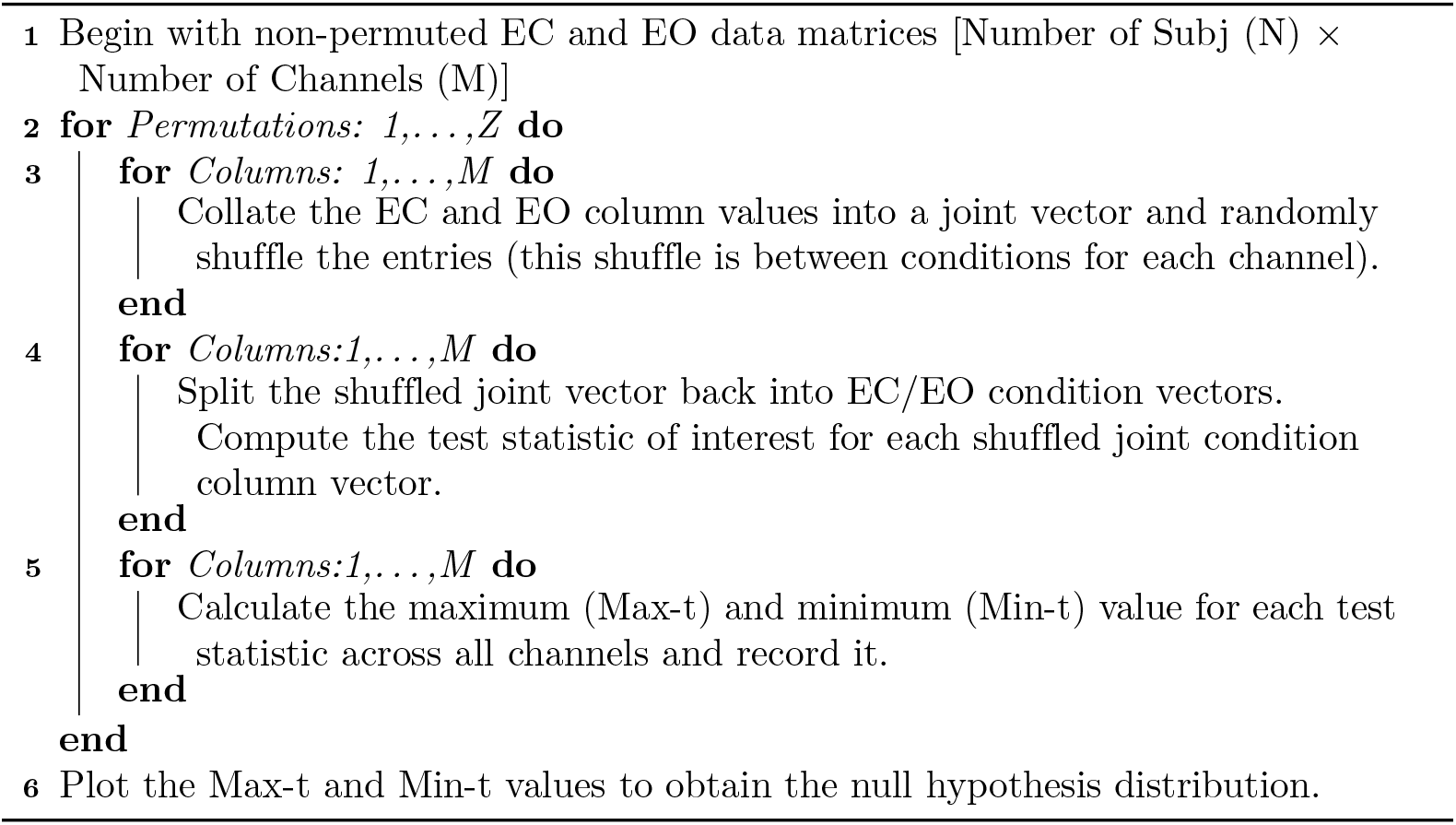

## Supporting information

**Fig. Area of** *cp*(*γ*) **between EC and EO conditions.** Empirical demonstration of the equivalence of the constant *c* in Eq 14 between EC and EO states.

**Appendix. Regularisation parameter selection.** Further details regarding the selection of the optimal regularisation parameter as well as an illustration of the systematic dependency of model fit on the regularisation parameter chosen.

**Appendix. Properties of the weakly damped measure (WDM).** Graphical illustration of the properties of the WDM defined in Eq 25.

## Acknowledgments

Many thanks to Prof. Suresh Muthukumaraswamy for providing access to the EC and EO resting state data used in Dataset 1. Special thanks to Prof. Conrad Perry and Prof. Peter Cadusch for their expertise and detailed feedback during the drafting process, it is greatly appreciated.

