## Supplementary material for "Alpha blocking and 1/*f^β^* spectral scaling in resting EEG can be accounted for by a sum of damped alpha band oscillatory processes": S1 Fig.

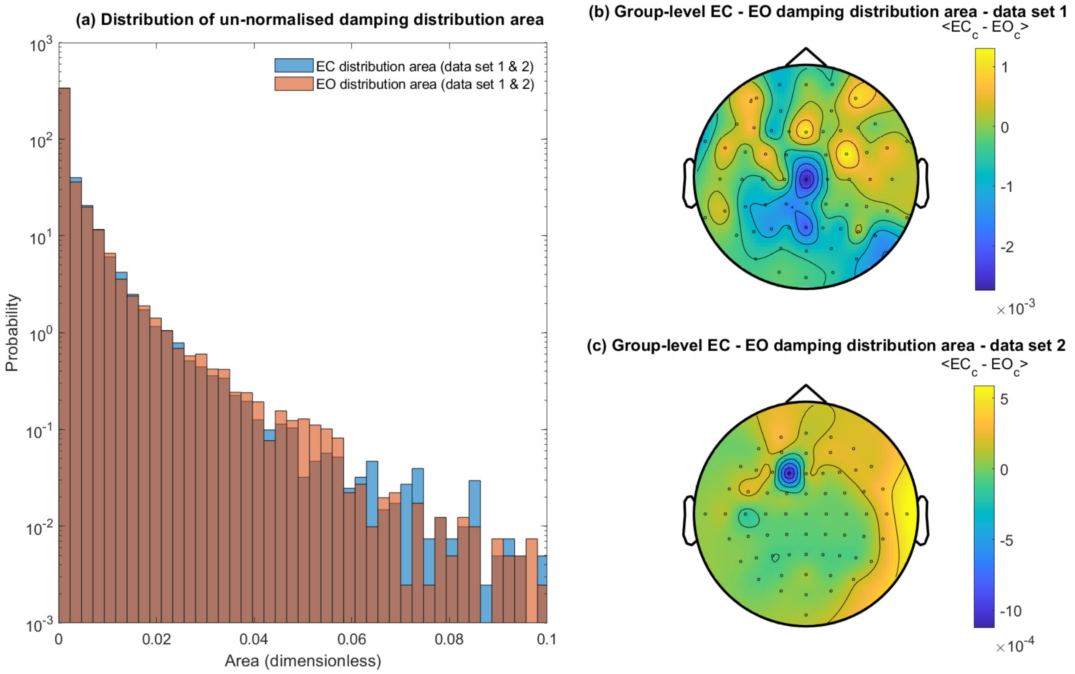


**Fig 1. EC and EO distribution and topogrpahic variation of the un-normalised area of the damping probability distribution ('c' constant).** **(a)** Pooled data analysis of the distribution of the constant 'c' in EC and EO states. The constant 'c' in Eq (14) can be calculated by finding the normalisation constant for the EC/EO damping distributions i.e. $1/A_{damp}$ where $A_{damp}$ is the non-normalised area of the estimated distribution of dampings. We note that there is no obvious difference in the distribution of the constant between resting states. **(b-c)** Topographic variation in the group-level difference in the constant between EC and EO states across data set 1 and 2. No statistically signifcant differences between states were found when using non-parametric permutation methods.
