## Supplementary material for "Alpha blocking and 1/*f^β^* spectral scaling in resting EEG can be accounted for by a sum of damped alpha band oscillatory processes": S2 Appendix

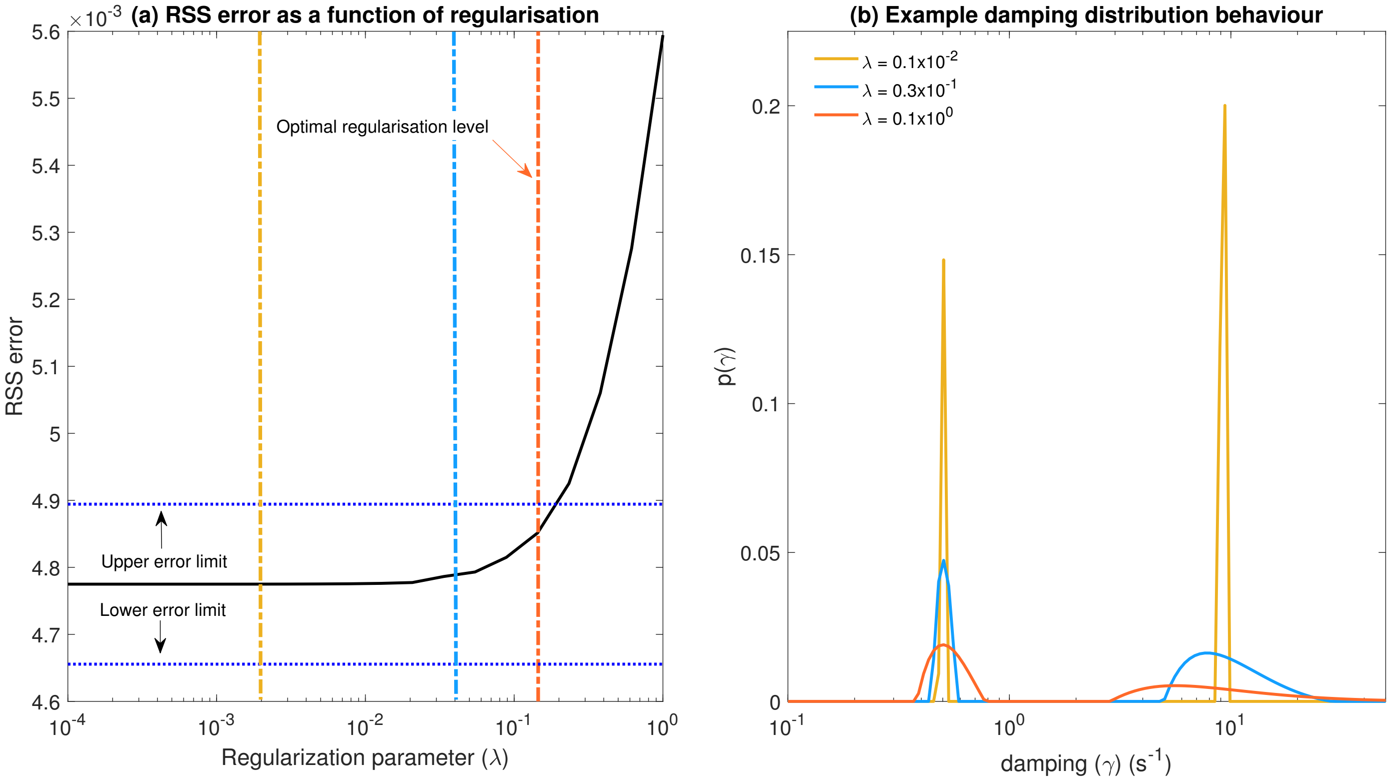


**Fig 1. Determining the optimal regularization parameter**. **(a)** Plot of the regularization parameter versus the RSS error between the measured and generated power spectrum with coloured (yellow, light blue, orange) vertical lines highlighting the specific regularisation values that correspond to the distributions found in the right panel. **(b)** Plots of the regularized damping distribution with increasing regularization parameter. The 'optimal' distribution is coloured orange and corresponds to the maximal entropy distribution that fell within the appropriate error bounds.

Regularization parameters were chosen on the basis of two criteria; choosing the maximum entropy inverse solution; minimizing the error between the data and model power spectrum. Fig 1a is presents the regularization parameter plotted against the RSS error for a participant with strong alpha blocking. In practice, we observe the error increasing with the magnitude of regularization. A plateau in the RSS is found where a reduction in the regularization parameter does not result in improved model fit. Optimal (best fit) distributions where chosen such that they had the maximum entropy but fell within approximately 2.5% of the minimum RSS. The orange vertical line in Fig 1a demonstrates the "optimal" regularization chosen for the particular example distribution.

In Fig 1b we observe the two distribution peaks respond in a systematic way to increases in the regularization parameter; amplitude of the modes decreases while increasing in width. The smoothing effect of the regularization is clearly observed as the distributions tend towards uniformity when significant regularization is present. The second distribution mode position varies to a greater extent than the weakly damped distribution modes position, and as such the second mode is more susceptible to changes in the regularization parameter. Irrespective of the behaviour present in the second distribution mode, the example behaviour shown in Fig 1b are consistent across sensors and subjects and thus provides confidence in the solutions obtained.
