## Supplementary material for "Alpha blocking and 1/*f^β^* spectral scaling in resting EEG can be accounted for by a sum of damped alpha band oscillatory processes": S3 Appendix

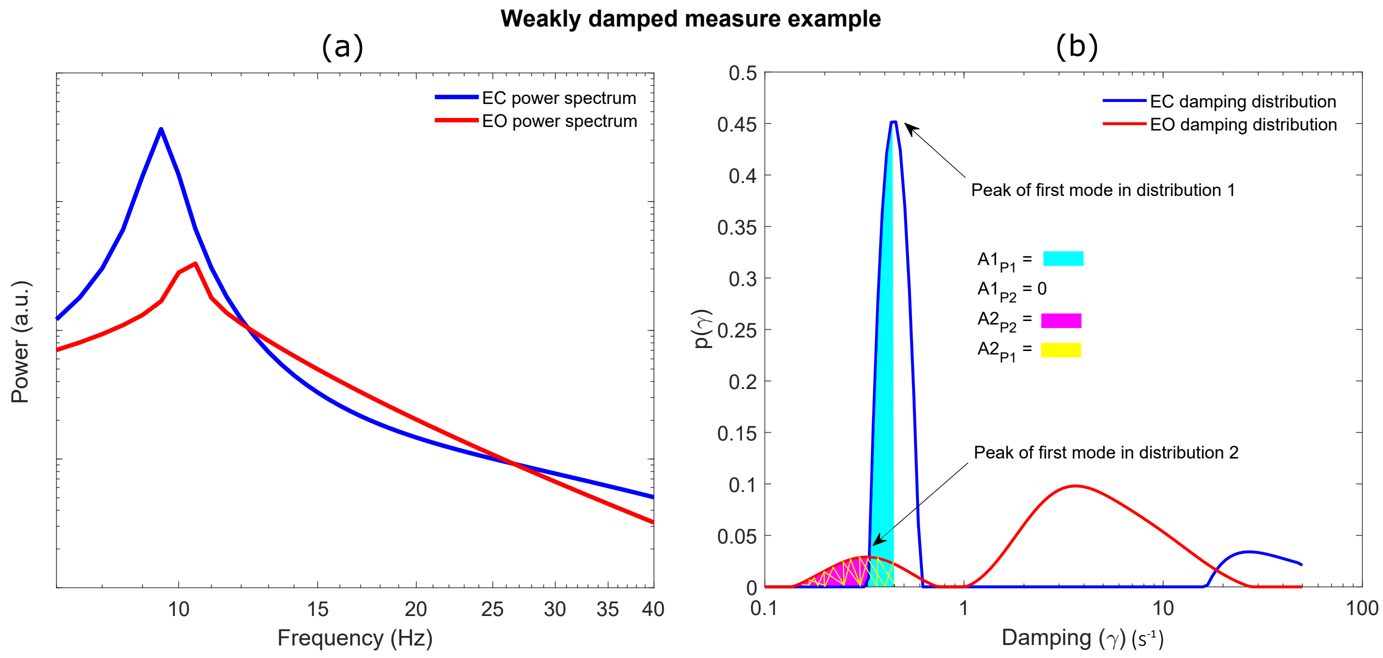


**Fig 1. Graphical exploration of weakly damped measure (a)** The EC and EO model fit power spectral densities computed using the respective damping distributions (right panel) in the forward model. **(b)** EC and EO damping probability distributions with the relevant distribution areas that go into the calculation of the weakly damped measure highlighted (light blue, magenta and yellow).

In this example we can clearly see that the alpha band activity in EC power spectrum is more weakly damped than the EO state. However, the distribution of dampings in the EO state peaks at a more weakly damped value that falls to the left of the first EC distribution mode. In this instance, if the weakly damped measure was calculated using only the $A1_{P2}$ and $A2_{P1}$ variables, then the measure for the EC state would be 0 as none of the area falls to the left of the EO peak. Whereas the EO state would have a value that is equal to the area highlighted with yellow, as it falls to the left of the EC peak. This would wrongly conclude that the EO state is more weakly damped than the EC. We can see that the including the area contained in the $A1_{P1}$ term (light blue), which is contributing to the larger alpha band activity, enables a fairer reflection of how weakly damped the distribution is in instances like that found in this example (note that the $A1_{P1}$ area is larger than the sum of the $A2_{P2}$ and $A2_{P1}$). The inclusion of the $A1_{P1}$ and $A2_{P2}$ terms corrects for these situations that may arise in either state and enables a more accurate reflection of which distribution is more weakly damped.
